# A novel method for multiple phenotype association studies based on genotype and phenotype network

**DOI:** 10.1101/2023.02.23.529687

**Authors:** Xuewei Cao, Shuanglin Zhang, Qiuying Sha

## Abstract

Joint analysis of multiple correlated phenotypes for genome-wide association studies (GWAS) can identify and interpret pleiotropic loci which are essential to understand pleiotropy in diseases and complex traits. Meanwhile, constructing a network based on associations between phenotypes and genotypes provides a new insight to analyze multiple phenotypes, which can explore whether phenotypes and genotypes might be related to each other at a higher level of cellular and organismal organization. In this paper, we first develop a bipartite signed network by linking phenotypes and genotypes into a Genotype and Phenotype Network (GPN). The GPN can be constructed by a mixture of quantitative and qualitative phenotypes and is applicable to binary phenotypes with extremely unbalanced case-control ratios in large-scale biobank datasets. We then apply a powerful community detection method to partition phenotypes into disjoint network modules based on GPN. Finally, we jointly test the association between multiple phenotypes in a network module and a single nucleotide polymorphism (SNP). Simulations and analyses of 72 complex traits in the UK Biobank show that multiple phenotype association tests based on network modules detected by GPN are much more powerful than those without considering network modules. The newly proposed GPN provides a new insight to investigate the genetic architecture among different types of phenotypes. Multiple phenotypes association studies based on GPN are improved by incorporating the genetic information into the phenotype clustering. Notably, it might broaden the understanding of genetic architecture that exists between diagnoses, genes, and pleiotropy.

## Introduction

Genome-wide association studies (GWAS) have successfully identified thousands of single nucleotide polymorphisms (SNPs) genetically associated with a wide range of complex human diseases and traits (Fine et al. 2019; Li et al. 2019). Over the past decade, more than 10,000 associations between SNPs and diseases/traits have been discovered (Visscher et al. 2017). Although GWAS have emerged as a common and powerful tool to detect the complexity of the genotype-phenotype associations, a common limitation of GWAS is that they focus on only a single phenotype at a time (Pendergrass et al. 2012; Pendergrass et al. 2013; Bush et al. 2016; Denny et al. 2016). Joint analysis of multiple correlated phenotypes for GWAS may provide more power to identify and interpret pleiotropic loci, which are essential to understand pleiotropy in diseases and complex traits (Bush et al. 2016; Verma et al. 2019; Lee et al. 2021). In brief, biological pleiotropy refers to a SNP or gene that has a direct biological influence on more than one phenotypic trait (Solovieff et al. 2013). Biological pleiotropy can offer significant insights in understanding the complex genotype-phenotype relationships (Li et al. 2019). Therefore, multiple phenotypes are usually collected in many GWAS cohorts and jointly analyzing multiple phenotypes may increase statistical power to discover the cross-phenotype associations and pleiotropy (Solovieff et al. 2013; Stephens 2013; Zhou and Stephens 2014; Sha et al. 2019).

Many statistical methods have been developed to jointly test the association between a SNP and multiple correlated phenotypes (Yang and Wang 2012). The most widely used methods for multiple phenotype association studies can be roughly classified into three categories: 1) statistical tests based on combining either the univariate test statistics or p-values, such as O’Brien’s method (O’Brien 1984), adaptive Fisher’s combination (AFC) (Liang et al. 2016), aSPU (Kim et al. 2015), and others (Yang et al. 2016); 2) multivariate analyses based on regression methods, such as multivariate analysis of variance (MANOVA) (Cole et al. 1994), reverse regression methods (MultiPhen) (O’Reilly et al. 2012), linear mixed effect models (LMM) (Laird and Ware 1982), and generalized estimating equations (GEE) (Liang and Zeger 1986); and 3) dimension reduction methods, such as clustering linear combination (CLC) (Sha et al. 2019), canonical correlation analysis (CCA) (Tang and Ferreira 2012), and principal components analysis (PCA) (Aschard et al. 2014; Wang et al. 2016). However, most phenotypes are influenced by many SNPs that act in concert to alter cellular function (Hawkins et al. 2010), the above mentioned methods are only based on phenotypic correlation without considering the genetic correlation among phenotypes. Therefore, these methods may loss statistical power to detect the true pleiotropic effects comparing the methods based on genetic architecture among complex diseases. To address this issue, numerous types of algorithms to investigate the genetic correlation among complex traits and diseases have been developed (Bulik-Sullivan et al. 2015; Pasaniuc and Price 2017; O’Connor and Price 2018). Many of these algorithms are often in conjunction with linkage disequilibrium (LD) information by using GWAS summary association data (Pasaniuc and Price 2017). For example, cross-trait LD score regression has been developed to estimate genetic and phenotypic correlation that requires only GWAS summary statistics and is not biased by overlapping samples (Bulik-Sullivan et al. 2015).

In 2007, a conceptually different approach based on the human disease network had been developed, exploring whether human complex traits and the corresponding genotypes might be related to each other at a higher level of cellular and organismal organization (Goh et al. 2007). Network analyses provide an integrative approach to characterize complex genomic associations (Gaynor et al. 2022). Therefore, constructing a network based on the associations between phenotypes and genotypes provides a new insight to simultaneously analyze multiple phenotypes and SNPs. Notably, it might broaden the understanding of genetic architecture that exists between diagnoses, genes, and pleiotropy (Verma et al. 2019). Modules detected from human disease networks are useful in providing insights pertaining to biological functionality (Tripathi et al. 2019). Therefore, community detection methods play a key role in understanding the global and local structures of disease interaction, in shedding light on association connections that may not be easily visible in the network topology (Newman 2018). Many community detection methods have been applied from social networks to human disease networks, such as Louvain’s method (Verma et al. 2019) with modularity as a measure and core module identification to identify small and structurally well-defined communities (Tripathi et al. 2019). However, most community detection methods have been developed for unsigned networks (Clauset et al. 2004; Newman and Girvan 2004; Barber 2007; Fortunato and Barthelemy 2007; Blondel et al. 2008; Newman 2012; Fortunato and Hric 2016).

To date, many biobanks, such as the UK Biobank (Sudlow et al. 2015), aggregate data across tens of thousands of phenotypes and provide a great opportunity to construct the human disease network and perform joint analyses of multiple correlated phenotypes. The electronic health record (EHR)-driven genomic research (EDGR) workflow is the most popular way to analyze multiple diagnosis codes in Biobank data, at its core, which is the use of EHR data for genomic research in the investigation of population-wide genomic characterization (Kohane 2011). In most EHR systems, the whole phenome can be divided into numerous phenotypic categories according to the first few characters of the International Classification of Disease (ICD) billing codes (Pendergrass and Crawford 2019). However, the ICD-based categories are based on the underlying cause of death rather than on the shared genetic architecture among all complex diseases and traits. Meanwhile, the phenotypes in large biobanks usually have extremely unbalanced case-control ratios. Therefore, linking phenotypes, especially EHR-derived phenotypes, with genotypes in a network is also very important to examine the genetic architecture of complex diseases and traits.

## Results

### Overview of Methods

In this paper, we develop a bipartite signed network by linking phenotypes and genotypes into a Genotype and Phenotype Network (GPN; **Figure 1a**). The GPN can be constructed by a mixture of quantitative and qualitative phenotypes and is applicable to phenotypes with extremely unbalanced case-control ratios for large-scale biobank datasets since the saddlepoint approximation (Dey et al. 2017) is used to test the association between genotype and phenotype with extremely unbalanced case-control ratio. After projecting genotypes into phenotypes, the genetic correlation of phenotypes can be calculated based on the shared associations among all genotypes (**Figure 1b**). We then apply a powerful community detection method to partition phenotypes into disjoint network modules using the hierarchical clustering method and the number of modules is determined by perturbation (**Figure 1c**) (Xie et al.). The phenotypes in each network module share the same genetic information. After partitioning phenotypes into disjoint network modules, a statistical method for multiple phenotype association studies can be applied to test the association between phenotypes in each module and a SNP, then a Bonferroni correction can be used to test if all phenotypes are associated with a SNP (**Figure 1d**). To jointly analyze the association between multiple phenotypes in each module with a SNP, we use six multiple phenotype association tests, including ceCLC (Wang et al. 2022), CLC (Sha et al. 2019), HCLC (Liang et al. 2022), MultiPhen (O’Reilly et al. 2012), O’Brien (O’Brien 1984), and Omnibus (Sha et al. 2019). The advantage of the association test based on network modules detected by GPN is that phenotypes in a network module are highly correlated based on the genetic architecture, therefore, the association test is more powerful to identify pleiotropic SNPs. After we obtain the GWAS signals from the previous steps, post-GWAS analyses can be applied to understand the high level of biological mechanism, such as pathway/tissue enrichment analysis and colocalization of GWAS signals and eQTL analysis in the specific disease-associated tissue (**Figure 1e-g**). The construction of GPN, community detection method, and six multiple phenotype association tests have been implemented in R, which is an open-source software and publicly available on GitHub: https://github.com/xueweic/GPN.

**Figure 1.**
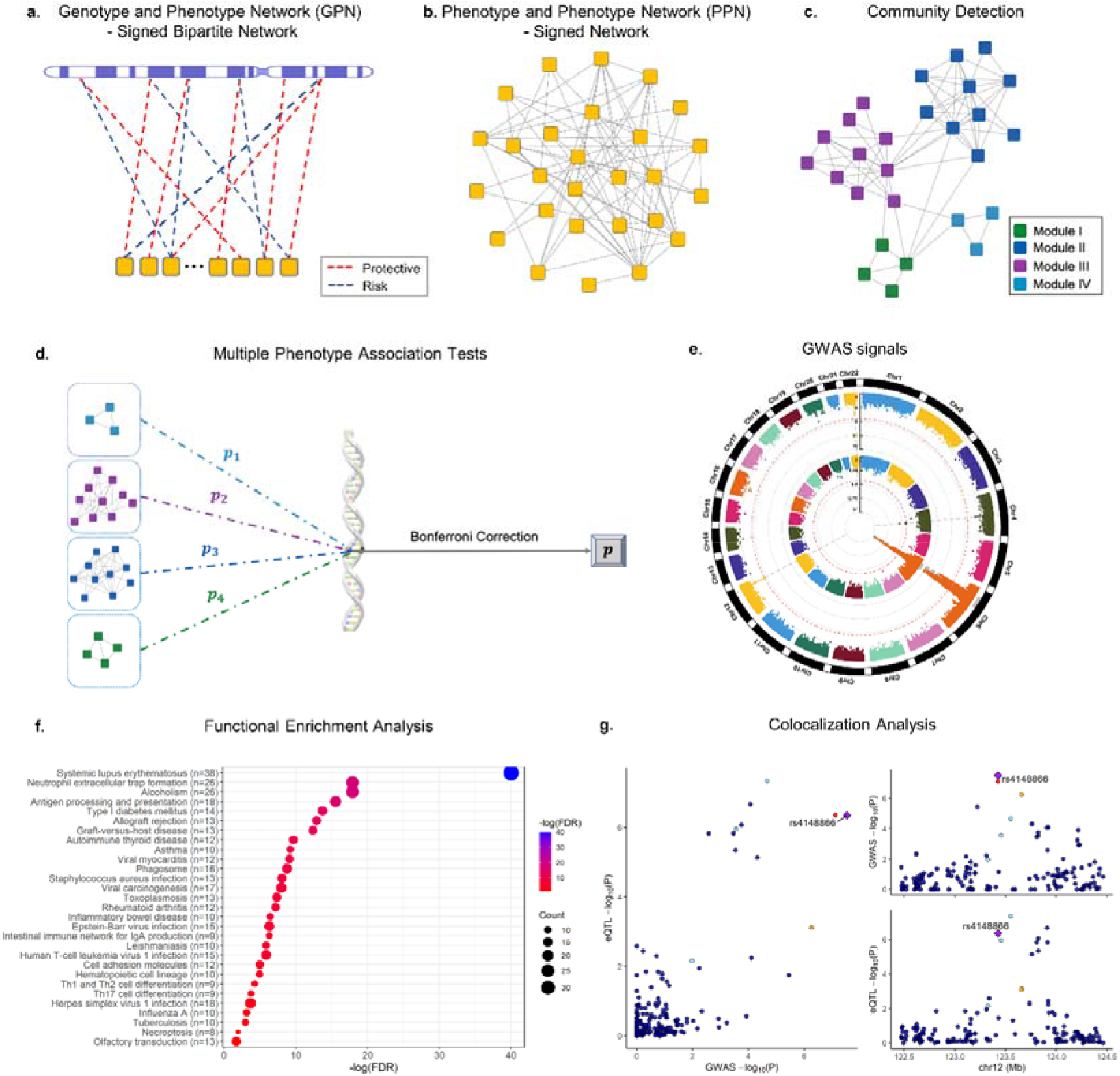
Overview of the method. **a**. Construction of GPN. Each phenotype (yellow square) and each SNP form a directed edge which represents the strength of the association, where the red dashed line indicates that the minor allele of the SNP is a protective allele to the phenotype, and the blue dashed line indicates that the minor allele of the SNP is a risk allele to the phenotype. **b**. Construction of PPN, which is the one-mode projection of GPN on phenotypes. **c**. The community detection method is used to partition phenotypes into disjoint network modules. **d**. Multiple phenotype association tests based on the network modules detected by GPN. **e**. GWAS signals are identified by a multiple phenotype association test with or without considering network modules. **f**. Functional enrichment analysis based on the detected GWAS signals and the publicly available functional database. **g**. Colocalization of GWAS signals and eQTL analysis.

### Simulation studies

We first use extensive simulation studies to validate multiple phenotype association studies based on the newly proposed GPN. In the simulation studies, we assess the type I error rate and power with different numbers of phenotypes (60, 80, and 100), different types of phenotypes along with different sample sizes: (i) mixture phenotypes are half quantitative and half qualitative with balanced case-control ratios for sample sizes of 2,000 and 4,000, and (ii) binary phenotypes are all qualitative but with extremely unbalanced case-control ratios for sample sizes of 10,000 and 20,000. Similar to the simulation models introduced in Sha et al. (Sha et al. 2019), we generate six different models (see Data Simulation).

#### Type I Error Rates

**Table S1-S6** summarize the estimated type I error rates of six multiple phenotype association tests for mixture phenotypes under models 1-6, respectively. “N.O.” represents the type I error rates of multiple phenotype association tests being calculated without considering network modules; “NET” presents the type I error rates of the tests being evaluated by considering network modules detected by GPN. Based on 500 Monte-Carlo (MC) runs which is the same as 10^6^ replicates, the 95% confidence intervals (CIs) for type I error rates divided by nominal significance levels 0.001 and 0.0001 are (0.938, 1.062) and (0.804,1.196), respectively. The bold-faced values indicate that the values are beyond the upper bounds of the 95% CIs. We can see that almost all of the estimated type I error rates of ceCLC, CLC, HCLC, and Omnibus tests are within 95% CIs. However, O’Brien in NET has inflated type I error rates under model 6. MultiPhen has inflated type I error rates for the sample size of 2,000. If the sample size is 4000, MultiPhen in N.O. also inflates type I error rates, but MultiPhen in NET can control type I error rates for the significance level is 0.0001. **Table S7-S12** summarize the estimated type I error rates of six multiple phenotype association tests for binary phenotypes with extremely unbalanced case-control ratios under models 1-6. Similar to **Tables S1-S6**, ceCLC, CLC, HCLC, and Omnibus have corrected type I error rates at almost all simulation settings. However, O’Brien in NET has inflated type I error rates and MultiPhen has inflated type I error rates at all scenarios.

#### Power comparisons

For power comparisons, we consider 100 causal SNPs for models 1-4 and 200 causal SNPs for models 5-6 (see Data Simulation). In each of the simulation models, the power is evaluated using 10 MC runs which is the same as 1,000 replicates for models 1-4 and 2,000 replicates for models 5-6. Meanwhile, the power is evaluated at the Bonferroni corrected significance level of 0.05 based on the number of causal SNPs in each MC run.

**Figure 2** (**Figure S1**) shows the power of six multiple phenotype association tests under six simulation models for different effect sizes with a total of 80 mixture phenotypes and a sample size of 4,000 (2,000). From **Figure 2** and **Figure S1**, we can see that: (i) All tests in NET (filled by the dashed line) are much more powerful than those in N.O., indicating that tests based on network modules detected by GPN are more powerful than the tests without considering network modules. Since the community detection method can partition phenotypes into different network modules based on shared genetic architecture, the phenotypes can be clustered in the same module if they have higher genetic correlations. In particular, the power of O’Brien (O’Brien 1984) increases a lot in the case of a SNP affecting phenotypes in different directions. (ii) ceCLC is more powerful than other tests in both N.O. and NET under the six simulation models. (iii) As sample size increases, the power of all multiple phenotype association tests increases. We also perform power comparisons for a total of 60 and 100 mixture phenotypes with 2,000 and 4,000 sample sizes for different effect sizes under the six simulation models (**Figures S2-S5**), respectively. We observe that the patterns of the power are similar to those observed in **Figure 2** and **Figure S1**.

**Figure 2.**
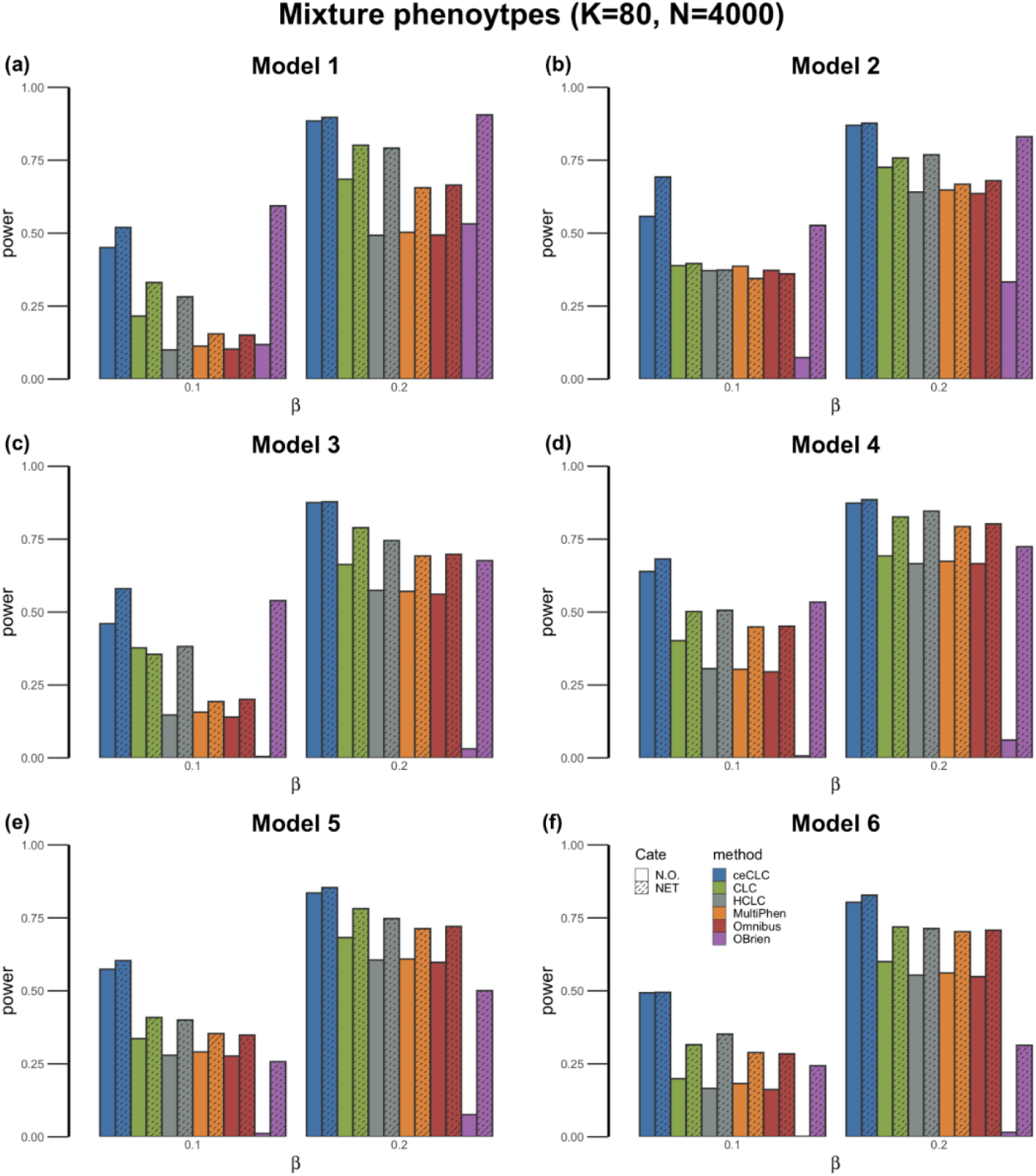
Power comparisons of the six tests as a function of effect size ^*β*^ under the six models. The number of mixture phenotypes (half continuous phenotypes and half binary phenotypes with balanced case-control ratios) is 80 and the sample size is 4,000. The power of all of the six tests is evaluated using 10 MC runs.

To mimic phenotypes in the UK Biobank, we also consider the case with all phenotypes being binary with extremely unbalanced case-control ratios. The phenotypes are generated based on extremely unbalanced case-control ratios which are randomly selected from the set of case-control ratios with cases greater than 200 from UK Biobank ICD-10 code level 3 phenotypes (see Real Dataset; case-control ratios belong to [0.000658,0.03937]). In this simulation, we consider a total of 60, 80, and 100 phenotypes along with two sample sizes, 10,000 and 20,000. **Figures S6-S11** show the power comparisons of the six tests under six simulation models. The patterns of power comparisons for binary phenotypes are similar to those observed in **Figure 2** and **Figure S1-S5**.

### Real Data Analysis based on UK Biobank

Furthermore, we apply the newly proposed multiple phenotype association test based on network modules detected by GPN to a set of diseases of the musculoskeletal system and connective tissue across more than 300,000 individuals from the UK Biobank.

#### Network Module Detection

We construct GPN based on 72 EHR-derived phenotypes in the diseases of the musculoskeletal system and connective tissue with 288,647 SNPs in autosomal chromosomes in the UK Biobank. Due to all phenotypes in our analysis being extremely unbalanced, the strength of the association between phenotype and genotype is calculated by the saddlepoint approximation (Dey et al. 2017). After the construction of GPN, we apply a powerful community detection method and these 72 phenotypes are partitioned into 8 disjoint network modules (**Figure 3**). There are 2-37 phenotypes in each module.

**Figure 3.**
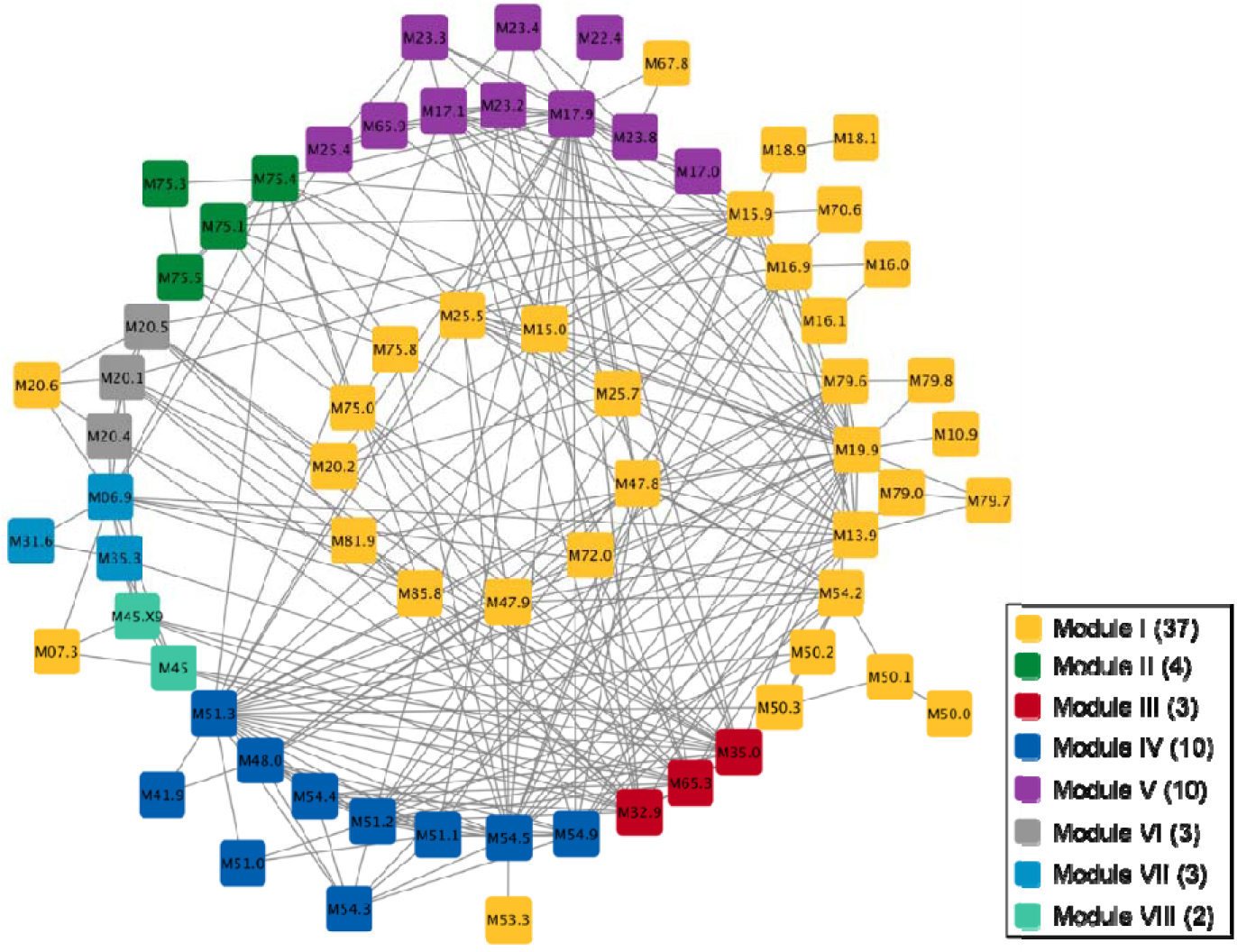
The network modules detected by the powerful community detection method based on GPN. The blocks with different color indicate different modules, where the values in the legend represent the number of phenotypes in each network module. The labels of phenotypes are listed in the form of ICD-10 code and the corresponding diseases can be found in the UK Biobank. The connection between two phenotypes represents the absolutely value of the weight greater than 40. The graph was prepared by Cytoscape.

We can see that the network modules are not consistent with the ICD-based categories which are based on the underlying cause of death rather than the shared genetic architecture among all complex diseases. For example, **Figure 3** shows three phenotypes, M32.9 Systemic lupus erythematosus, M35.0 Sicca syndrome, and M65.3 Trigger finger, are detected in network module III (in red). However, these three phenotypes do not belong to the same ICD-category (Data-Field 41202 in UK Biobank), where M35.0 is one of the diseases in the other systemic involvement of connective tissue (M35) and M65.3 belongs to the synovitis and tenosynovitis (M65). To investigate the genetic correlation among these three phenotypes, we use the saddlepoint approximation to test the association between each phenotype and each SNP. As shown in **Figure S12**, the Manhattan plots for the three phenotypes in network module III (M32.9, M35.0, and M65.3) have a similar pattern. Although the synovitis and tenosynovitis (M65.9) and M65.3 belong to the same ICD code category (M65), the Manhattan plot of M65.9 shows that there are no SNPs significantly associated with this phenotype and the genetic correlation between M65.9 and M65.3 is not strong. Therefore, we can conclude that the community detection method based on our proposed GPN can partition phenotypes into different categories based on the shared genetic architecture.

Furthermore, we apply the hierarchical clustering method to compare the genetic correlation of phenotypes obtained by our proposed GPN and that estimated by LDSC (Bulik-Sullivan et al. 2015). **Figures S13-S14** show that dendrograms of hierarchical clustering method based on the genetic correlation of phenotypes obtained by GPN, and the phenotypic or genetic correlation estimated by LDSC, respectively. In **Figure S13**, the cluster results of the phenotypic correlation estimated by LDSC are similar to that of the genetic correlation based on GPN, but GPN can separately identify two highly genetic correlated phenotypes, ankylosing spondylitis (M45) and ankylosing spondylitis with site unspecified (M45.X9). However, the cluster results of the genetic correlation estimated by LDSC are different from those obtained by GPN. Some phenotypes in the same UK Biobank level 1 category can be clustered in the same group by GPN but not by LDSC (**Figure S14**).

#### Interpretation of the Association Test

We apply five multiple phenotype association tests (ceCLC, CLC, HCLC, O’Brien, and Omnibus) to test the association between 72 EHR-derived phenotypes and each of 288,647 SNPs in the UK Biobank. MultiPhen is not considered here since it has inflated type I error rates, especially for the phenotypes with extremely unbalanced case-control ratios.

First, we apply the five tests in N.O. to test the association between 72 phenotypes and each SNP. We use the commonly used genome-wide significance level 5×10^-8^. **Figure 4(a)** shows the Venn diagram of the number of SNPs identified by the five tests. There are 11 SNPs identified by all five tests. ceCLC identifies 647 SNPs with 32 unique SNPs not being identified by other four tests. Among the 32 novel SNPs, two SNPs, rs13107325 (p-value = 4.6×10^-10^) and rs443198 (p-value = 1.73×10^-11^), are significantly associated with at least one of the 72 phenotypes reported in the GWAS catalog (**Table S14**). rs13107325 is reported to be associated with osteoarthritis (M19.9) (Tachmazidou et al. 2019) and rotator cuff syndrome (M75.1) (Kim et al. 2021). Meanwhile, rs13107325 is mapped to gene *SLC39A8* that is also reported to be significantly associated with multisite chronic pain (M25.5) (Johnston et al. 2019). rs443198 is mapped to gene *NOTCH4* which is associated with systemic sclerosis (M34) (Gorlova et al. 2011). Moreover, the mapped gene *NOTCH4* is one of the most important genes reported to be associated with multiple diseases in the disease category of the musculoskeletal system and connective tissue, such as rheumatoid arthritis (M06.9) (Terao et al. 2011), psoriatic arthritis (M07.3) (Aterido et al. 2019), Takayasu arteritis (M31.4) (Renauer et al. 2015), systemic lupus erythematosus (M32.9) (Chung et al. 2014), and appendicular lean mass (M62.9) (Cordero et al. 2019). We map these 32 unique SNPs into genes with 20 kb upstream and 20 kb downstream regions. There are 27 out of 32 SNPs with corresponding mapped genes associated with 14 phenotypes reported in the GWAS catalog (**Table S14**). These 14 phenotypes and corresponding ICD-10 codes are summarized in **Table S15**.

**Figure 4.**
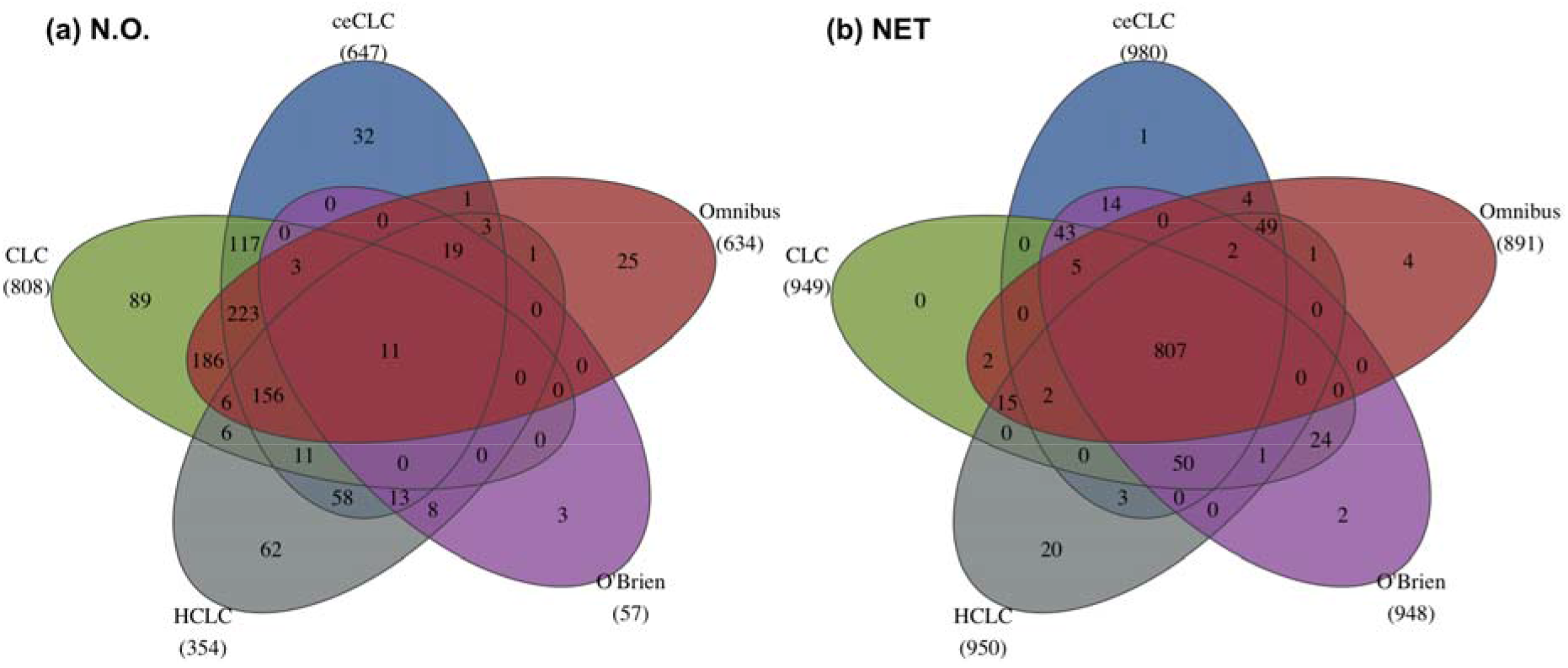
The Venn diagram of the number of SNPs identified by ceCLC, CLC, HCLC, O’Brien, and Omnibus in N.O. (**a**) and in NET (**b**). The number below each method indicates the total number of SNPs identified by the corresponding method.

Next, we test the associations between phenotypes in each of the eight network modules detected by the GPN and each SNP. Then, we adjust the p-value of each method for testing the association between a SNP and all of the 72 phenotypes by Bonferroni correction. We adopt the commonly used genome-wide significance level 5×10^-8^. **Figure 4(b)** shows that all tests can identify more SNPs comparing with the number of SNPs identified in N.O. ceCLC in NET identifies 980 SNPs, where 647 SNPs are identified in N.O. Meanwhile, there are 950 SNPs identified by HCLC, 949 SNPs by CLC, and 891 SNPs by Omnibus, where the corresponding results in N.O. are 354 SNPs, 808 SNPs, and 634 SNPs, respectively. In particular, the number of SNPs identified by O’Brien in NET is increased a lot, where there are 948 SNPs identified in NET and only 57 SNPs identified in N.O. As the results shown in **Figure 4(b)**, there are 807 overlapped SNPs identified by all five tests in NET which is much larger than 11 overlapped SNPs identified in N.O.

To compare the difference between the tests in N.O. and in NET, we summarize the number of overlapping SNPs identified by each method in N.O. and NET in **Figure S15**. We observe that most SNPs identified in N.O. can be identified in NET. Meanwhile, tests in NET can identify much more SNPs than those in N.O. As mentioned previously, the advantage of the tests based on the network modules detected by GPN is that we can identify potential pleiotropic SNPs and also interpret SNP effects on which network modules based on the shared genetic architecture. Notably, we also investigate the smallest p-value obtained by each of the eight phenotypic modules for each of the 980 SNPs identified by ceCLC. For example, 396 SNPs have the smallest p-values for testing the association with network module III. Based on the results of the univariate score test corrected for saddlepoint approximation (SPAtest) (**Figure S12**), 104 SNPs are significantly associated with at least one phenotype in module III. All of these 104 SNPs can be identified by ceCLC, HCLC, and Omnibus in NET and 103 SNPs can be identified by CLC and O’Brien in NET. The results show that the tests based on network modules can detect potential pleiotropic loci which can not be detected by the univariate test.

#### Pathway Enrichment Analysis

ceCLC is more powerful than the other four tests in simulations and also can identify more SNPs in real data analysis, therefore, we only perform the post-GWAS analyses of the SNPs identified by ceCLC. There are 191 mapped genes containing at least one of the 647 SNPs identified by ceCLC in N.O. and 252 mapped genes containing at least one of the 980 SNPs identified by ceCLC in NET. In this study, significantly enriched pathways are identified by those genes with false discovery rate (FDR) < 0.05.

From the pathway enrichment analyses, we observe that ceCLC based on the network modules identifies more significantly enriched pathways than that without considering network modules. **Figure 5** shows that 16 pathways are significantly enriched by 191 mapped genes in N.O. and 29 pathways are significantly enriched by 252 mapped genes in NET, where all of the 16 pathways identified in N.O. are also identified in NET. Two pathways identified in N.O. and NET, rheumatoid arthritis (hsa05323; FDR = 8.72×10^-3^ in N.O. and FDR = 6.48×10^-8^ in NET) and systemic lupus erythematosus (hsa05322; FDR = 4.25×10^-19^ in N.O. and FDR = 1.02×10^-40^ in NET) showed in **Figure 5**, are related to the diseases of the musculoskeletal system and connective tissue. For example, osteopetrosis (M19.9) and rheumatoid arthritis (M06.9) are related to the rheumatoid arthritis pathway. Meanwhile, the pathway related to at least one of the 72 phenotypes, hematopoietic cell lineage (hsa04640; FDR = 1.08×10^-5^), is only identified in NET. Notably, DBGET system (https://www.genome.jp/dbget-bin/www_bget?hsa05322) reports that there are two pathways related to systemic lupus erythematosus: antigen processing and presentation (hsa04612; FDR = 4.83×10^-3^ in N.O. and FDR = 2.82×10^-16^ in NET) identified in both N.O. and NET and cell adhesion molecule (hsa04514; FDR = 1.04×10^-5^) only identified in NET.

**Figure 5.**
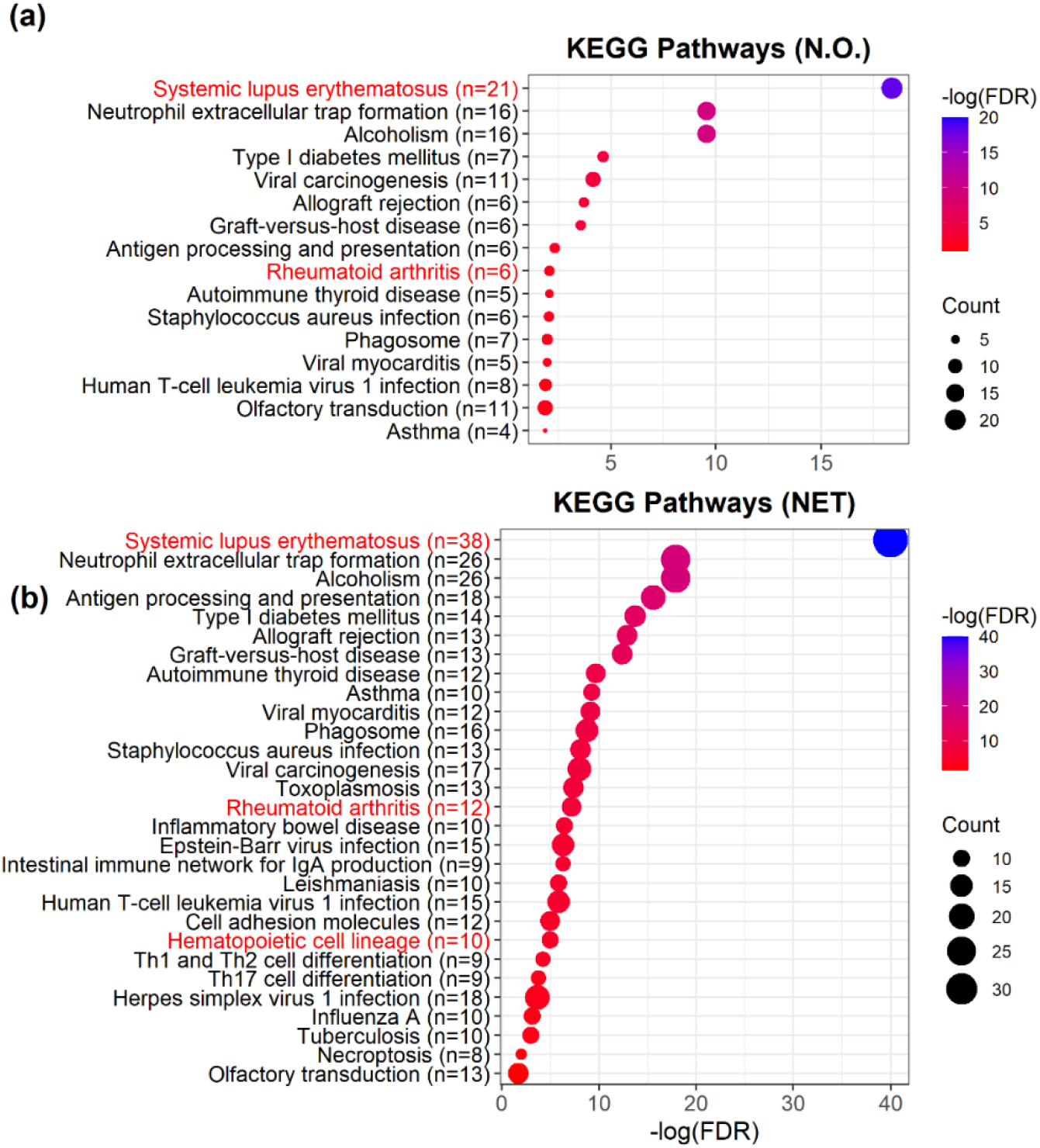
The results for the pathway enrichment analysis based on the genes identified by ceCLC and the KEGG database in N.O. (**a**) and NET (**b**). The red marked pathways denote the pathways related to the diseases of the musculoskeletal system and connective tissue. There are 191 genes in N.O. and 252 genes in NET that are applied to the pathway enrichment analysis.

Meanwhile, the above five pathways related to the diseases of the musculoskeletal system and connective tissue contain more enriched genes identified by ceCLC in NET than the enriched genes identified in N.O. For example, 43 SNPs within six mapped genes identified by ceCLC in N.O. are enriched in rheumatoid arthritis pathway, including *ATP6V1G2, HLA-DRA, LTB, TNF, HLA-DRB1*, and *HLA-DQA1*; and 111 SNPs within 12 mapped genes in NET are enriched in this pathway, including *HLA-DMA, HLA-DMB, ATP6V1G2, HLA-DRA, LTB, HLA-DOA, TNF, HLA-DOB, HLA-DQA2, HLA-DRB1, HLA-DQA1*, and *HLA-DQB1*. Compared with the results of ceCLC in N.O., the test based on network modules identifies six more enriched genes, especially, gene *HLA-DMB* (including rs241458; p-value = 7.09×10^-9^) and gene *HLA-DOA* (including rs3097646; p-value = 5.50×10^-9^) that have not been reported in the GWAS catalog.

#### Tissue Enrichment Analysis

To further investigate the biological mechanism, we use FUMA (Watanabe et al. 2017) to annotate 191 mapped genes in N.O. and 252 mapped genes in NET in terms of biological context. Due to these mapped genes associated with at least one phenotype in the diseases of the musculoskeletal system and connective tissue, we can test if these mapped genes are enriched in the relevant-tissue based on FUMA. **Figure S16** shows the ordered enriched tissues based on the mapped genes identified by ceCLC in N.O. and NET. We observe that the mapped genes identified by ceCLC in N.O. are most enriched in brain-related tissue (**Figure S16(a)**). Nevertheless, **Figure S16(b)** shows that the mapped genes identified by ceCLC in NET are significantly enriched in the Muscle-Skeletal tissue with p-value < 0.05. The construction of GPN is benefit to multiple phenotype association studies by clustering the related phenotypes based on the genetic information. Notably, the identified SNPs are more likely to be within the same relevant biological context.

#### Colocalization of GMAS and eQTL analysis

We perform the colocalization analysis on the 33 unique SNPs identified by ceCLC (**Table S13**; one SNP in NET and 32 SNPs in N.O.) and all SNP-gene association pairs in the Muscle Skeletal tissue reported in GTEx. **Figure S17** shows the colocalization signals with the uniquely identified SNPs by ceCLC that are selected to be the lead SNPs in the colocalization analysis. NET identifies one unique SNP, rs4148866, which is mapped to gene *ABCB9*. Even if gene *ABCB9* has no reported associations with any diseases of the musculoskeletal system and connective tissue in the GWAS Catalog, the Bayesian posterior probability of colocalization analysis for shared variant of significant SNPs identified by ceCLC and gene expression in the Muscle Skeletal tissue (PP_H4_) is 98.4%. The higher value of PP_H4_ indicates that gene *ABCB9* and Muscle Skeletal tissue play an important role in the disease mechanism due to the same variant responsible for a GWAS locus and also affecting gene expression (Hormozdiari et al. 2016). Among 32 unique SNPs identified by ceCLC in N.O., there are two SNPs, rs34333163 and rs6916921, selected to be the lead SNPs (**Figure S17**). Both of them are reported in the GWAS Catalog that have associations with at least one of the diseases in the musculoskeletal system. However, the PP_H4_ values for the corresponding genes *SLC38A8* and *ATP6V1G2* are lower than 50%.

## Discussion

In this paper, we propose a novel method for multiple phenotype association studies based on genotype and phenotype network. The construction of a bipartite signed network, GPN, is to link genotypes with phenotypes using the evidence of associations. To understand pleiotropy in diseases and complex traits and explore the genetic correlation among phenotypes, we project genotypes into phenotypes based on the GPN. We also apply a powerful community detection method to detect the network modules based on the shared genetic architecture. In contrast to previous community detection methods for disease networks, the applied method benefits from exploring the biological functionality interactions of diseases based on the signed network. Furthermore, we apply several multiple phenotype association tests to test the association between phenotypes in each network module and a SNP. Extensive simulation studies show that all multiple phenotype association tests based on network modules have corrected type I error rates if the corresponding test is a valid test for testing the association between a SNP and phenotypes without considering network modules. Most tests in NET are much more powerful than those in N.O. Meanwhile, we evaluate the performance of the association tests based on network modules detected by GPN through a set of 72 EHR-derived phenotypes in the diseases of the musculoskeletal system and connective tissue across more than 300,000 samples from the UK Biobank. Compared with the tests in N.O., all tests based on network modules can identify more potentially pleiotropic SNPs and ceCLC can identify more SNPs than other methods.

In addition, the construction of GPN does not require access to individual-level genotypes and phenotypes data, which only requires association evidence between each genotype and each phenotype. Therefore, when individual-level data are not available, this evidence can be obtained from GWAS summary statistics, such as the effect sizes (odds ratios for binary phenotypes) and corresponding p-values. Meanwhile, the simulation studies show that the powerful network community detection method can correctly partition phenotypes into several disjoint network modules based on the shared genetic architecture. Since the determination of the number of network modules in community detection method is independent of the association tests (Xie et al.), we only need to perform the perturbation procedure once in real data analyses. In our real data analysis with 72 phenotypes and 288,647 SNPs, it only takes 1.5 hour with 1,000 perturbations to obtain the optimal number of network modules on a macOS (2.7 GHz Quad-Core Intel Core i7, 16 GB memory).

In summary, the proposed GPN provides a new insight to investigate the genetic correlation among phenotypes. Especially when the phenotypes have extremely unbalanced case-control ratios, the weight of an edge in the signed bipartite network can be calculated based on the saddlepoint approximation. The power of multiple phenotype association tests based on network modules detected by GPN are improved by incorporating the genetic information into the phenotypic clustering. Therefore, the proposed method can be applied to large-scale data across multiple related traits and diseases (i.e., biobanks data set, etc.).

## Methods

Consider a sample with *n* unrelated individuals, indexed by *i* = 1,···, *n*. Suppose each individual has a total of *K* phenotypes and *M* SNPs. Let **Y** = (*y_ik_*) be an *n×K* matrix of *K* phenotypes, where *y_ik_* denotes the phenotype value of the *i^th^* individual for the *k^th^* phenotype. The phenotypes can be both quantitative and qualitative, especially for phenotypes with extremely unbalanced case-control ratios. Let **G** = (*g_im_*) be an *n×M* matrix of genotypes, where *g_im_* represents the genotypic score of the *i^th^* individual at the *m^th^* SNP which is the number of minor alleles that the *i^th^* individual carries at the SNP.

### Construction of the Genotype and Phenotype Network

We first introduce a signed bipartite genotype and phenotype network (GPN) (**Figure 1a**). The weight of an edge represents the strength of the association between the two nodes (one is the phenotype and the other one is the genotype). The strength of the association has two directions, positive and negative. The adjacency matrix of GPN is a *K×M* matrix **T** = (*T_km_*), where *T_km_* represents the strength of the association between the *k^th^* phenotype and the *m^th^* SNP. To calculate the adjacency matrix **T**, we consider both the strengths and the directions of the associations. We first consider that there are no covariates. The strength of the association *T_km_* can be estimated by the score test statistic 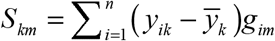 and its p-value *P_km_* under the generalized linear models *g*(*E*(*Y_ik_|g_im_*)) = *β*_0*km*_ + *β*_1*km*_*g*_*im*_ (*k* = 1,…, *K* and *m* = 1,···, *M*) (Sha et al. 2011). Here, 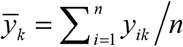 and *g*() is a monotonic link function. Two commonly used link functions are the identity link for quantitative traits and the logit link for binary traits. If there are *p* covariates for the *i^th^* individual, *x*_*i*1_,···, *x_ip_*, we adjust genotype and phenotype for the covariates using the following linear models proposed by Price et al. (Price et al. 2006) and Sha et al. (Sha et al. 2012),

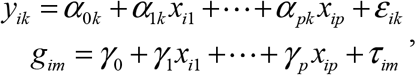

where *ε_k_* = (*ε*_1*k*_,···,*ε_nk_*)^*T*^ and ***τ**_m_* = (*τ*_1*m*_,···, *τ_nm_*)*^T^* denote the error terms of the *k^th^* phenotype and the *m^th^* SNP, respectively. We use the residuals of the respective linear model to replace the original genotypes and phenotypes.

For quantitative traits or binary traits with fairly balanced case-control ratios, we can use the normal approximation of 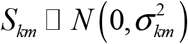 to calculate p-value *p_km_* under the null hypothesis that the *k^th^* phenotype and the *m^th^* SNP have no association, where 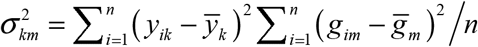 and 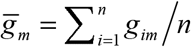. Dey et (Dey et 2017) pointed out that a normal approximation of *S_km_* has inflated type I error rates for binary traits with unbalanced case-control ratios. Therefore, we use saddlepoint approximation to calculate the p-value *p_km_* for the phenotypes with unbalanced, especially extremely unbalanced case-control ratios (Dey et al. 2017). We define the (*k,m*)^*th*^ element of the adjacency matrix of GPN, *T_km_*, as 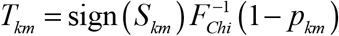, where *F_chi_* ( ) denotes the CDF of 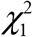 That is, we use sign(*S_km_*) to define the direction of the association and use 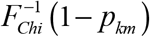 to define the strength of the association. *T_km_* > 0 and *T_km_* < 0 represent two directions of the association between the *k^th^* phenotype and the *m^th^* SNP. If *T_km_* > 0, the minor allele of the *m^th^* SNP is a protective allele to the *k^th^* phenotype; if *T_km_* < 0, the minor allele of the *m^th^* SNP is a risk allele to the *k^th^* phenotype.

Although a bipartite network may give the most complete representation of a particular network, it is often convenient to work with just one type of nodes, that is, phenotypes or genotypes. The Phenotype and Phenotype Network (PPN) is the one-mode projection of GPN on phenotypes. In PPN, nodes only represent phenotypes (**Figure 1b**). Let **W** = (*W_k1_*) denote the adjacency matrix of the PPN in which each edge has a positive or negative weight. We define *W_k1_* as the weight of the edge connecting the *k^th^* and *l^th^* phenotypes, which is given by

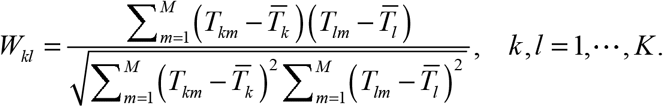

Here, *W_k1_* is the genetic correlation between the *k^th^* and *l^th^* phenotypes based on the association strengths *T_km_* for *k* = 1,···, *K* and *m* = 1,···, *M*. Thus, the PPN is also a signed network.

### Community Detection Method

We apply a powerful community detection method to partition *K* phenotypes into disjoint network modules using the Ward hierarchical clustering method with a similarity matrix defined by the genetic correlation matrix **W**(Xie et al.). The number of network modules is determined by the following perturbation procedure (Nguyen et al. 2017). In details, we first use the Ward hierarchical clustering method to group the *K* phenotypes into *k*_0_ (*k*_0_ = 1,···, *K*–1) clusters and build the *K × K* connectivity matrix **C**_*k*_0__ with the (*k,1*)^*th*^ element of matrix **C**_*k*_0__ given by

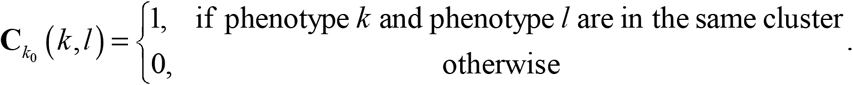

Then, we generate *B* perturbed data sets. The *b^th^* perturbed data set is generated by 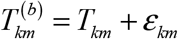, where *ε_km_* □ *N*(0, *σ*^2^), *σ*^2^ = median(Var (***T***_1_),···, Var (***T**_M_*)), and ***T**_m_* = (*T_1m_*,···, *T_Km_*). We denote the connectivity matrix of *k*_0_ cluster based on the *b^th^* perturbed data set by 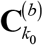. Let 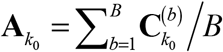 and **D**_*k*_0__ = |**A**_*k*_0__ – **C**_*k*_0__|, *F*_*k*_0__ denotes the empirical CDF of the elements of **D**_*k*_0__, and *AF*_*k*_0__ denotes the area under the curve of *F*_*k*_0__, where *F*_*k*_0__ (*x*) = #{**D**_*k*_0__ (*1,k*) ≤ *x*: *1*, *k* = 1,···, *K*}/*K*^2^. Then, the optimal number of network modules is given by

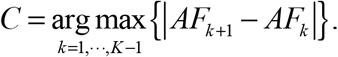

We can use the identified *C* network modules to further investigate the associations between phenotypes in each network module and SNPs.

### Multiple Phenotype Association Tests

After we obtain *C* network modules for the phenotypes, we apply a multiple phenotype association test to identify the association between phenotypes in each of the *C* network modules and a SNP. Any multiple phenotype association test can be applied here. In this article, we apply six commonly used multiple phenotype association tests to each network module, including ceCLC (Wang et al. 2022), CLC (Sha et al. 2019), HCLC (Liang et al. 2022), MultiPhen (O’Reilly et al. 2012), O’Brien (O’Brien 1984), and Omnibus (Sha et al. 2019) (see details in **Text S1**), then a Bonferroni correction is used to adjust for multiple testing for the *C* network modules to test if all phenotypes in the *C* network modules associated with a SNP.

### Data Simulation

We conduct comprehensive simulation studies to evaluate the type I error rates and powers of multiple phenotype association tests based on network modules detected by GPN and compare them to the powers of the corresponding tests without considering network modules. To evaluate the performance of our proposed method, we consider different types of phenotypes: (i) mixture phenotypes: half quantitative and half qualitative with balanced case-control ratios, and (ii) binary phenotypes: all qualitative but with extremely unbalanced case-control ratios. We generate *N* individuals with *M* SNPs and *K* phenotypes. The genotypes at *M* SNPs are generated according to the minor allele frequency (MAF) under Hardy-Weinberg Equilibrium (HWE). Below, we first describe how to generate quantitative phenotypes. Suppose that there are *C* phenotypic categories and *k* = *K/C* phenotypes in each phenotypic category. Let **Y**_*c*_ = (***y**_c1_*,···, ***y**_ck_*) denote the phenotypes in the *c^th^* category. Similar to Sha et al. (Sha et al. 2019), we generate *k* quantitative phenotypes in each category using the following factor model,

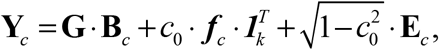

where **G** = (***G***_1_,···, ***G**_M_*) is the matrix of *M* SNPs with dimension *N×M* which are generated from a binomial(2, *MAF*) distribution for each SNP; **B**_*c*_ is an *M* × *k* matrix of effect sizes of *M* SNPs on *k* phenotypes in the *c^th^* phenotypic category; **E**_*c*_ □ *MVN_k_* (***0*, Σ**) is an *N*×*k* matrix of error term with **Σ** = (*σ_ij_*), where *σ_ij_* = *ρ*^|*i−j*|^ and *ρ* is a constant between 0 to 1; ***f**_c_* is a factor vector in **f** = (***f***_1_,··· ***f**_C_*) which follows *MVN_C_*(***0*, Σ**_*f*_), where **Σ**_*f*_ = (1–*ρ_f_*)**I***_C_* + *ρ_f_***J**_*C*_, *ρ_f_* =corr (***f**_i_*, ***f**_j_*) if *i* ≠ *j*, **J**_*C*_ is a *C*×*C* matrix with all elements of 1, and **I**_*c*_ is the identity matrix; *c_0_* is a constant number which represents a proportion. Therefore, the correlation between the *i^th^* phenotype and the *j^th^* phenotype within each category is 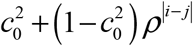 and the between-category correlation is 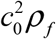.

To generate a qualitative disease affection status, we use a liability threshold model based on a quantitative phenotype and its case-control ratio. Let *n_a_* and *n_c_* denote the number of affected individuals and the number of non-affected individuals. For a given casecontrol ratio *r* and sample size *N*, *n_c_* = *N/*(*r*+1) and *n_a_* = *rN/*(*r* +1). An individual is defined to be affected if the individual’s phenotype is in the top *n_a_* of all phenotypes. For each phenotype, the case-control ratio is randomly chosen from a set *S*. The set *S* contains all case-control ratios with the number of cases greater than 200 from UK Biobank ICD-10 code level 3 phenotypes.

Based on the factor model, we consider different numbers of phenotypes, 60, 80, and 100, and different sample sizes. For mixture phenotypes, the sample sizes are 2,000 and 4,000; for binary phenotypes, the sample sizes are 10,000 and 20,000. We consider six simulation models (**Text S2** and **Table S15**) with *M* = 2,000, *MAF □ U*(0.05,0.5), *ρ* = 0.3, 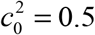, and 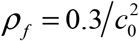 (between-category correlation is 0.3). The calculations of the type I error rates and power of multiple phenotype association test in N.O. and in NET are summarized in **Text S3**.

### Real Dataset

The UK Biobank is a population-based cohort study with a wide variety of genetic and phenotypic information (Bycroft et al. 2018). It includes ~ 500K people from all around the United Kingdom who were aged between 40 and 69 when recruited in 2006-2010 (Sudlow et al. 2015; Bycroft et al. 2017). Following the genotype and phenotype preprocess introduced in Liang et al. (Liang et al. 2022), there are 288,647 SNPs and 72 EHR-derived phenotypes in the diseases of the musculoskeletal system and connective tissue for 322,607 individuals are kept in our real data analysis (**Text S4** and **Figure S18**). Among the 72 phenotypes, lumbar and other intervertebral disk disorders with myelopathy (M51.0) has the smallest case-control ratio 0.000658 with 212 cases and 322,395 controls; Gonarthrosis (M17.9) has the largest case-control ratio 0.03937 with 12,218 cases and 310,389 controls. Therefore, all of the phenotypes we considered in our analysis have extremely unbalanced case-control ratios. Furthermore, each phenotype is adjusted by 13 covariates, including age, sex, genotyping array, and the first 10 genetic principal components (PCs) (Sha et al. 2012). The analysis is performed based on the adjusted phenotypes.

### Correlation Analysis

To compare the genetic and phenotypic correlations among the 72 EHR-derived phenotypes, we apply cross-triat LDSC regression (Bulik-Sullivan et al. 2015) to obtain the genetic correlation and phenotypic correlation which can provide useful etiological insights (Bulik-Sullivan et al. 2015). GWAS summary statistics are generated from the association between phenotype and genotype which are calculated by the saddlepoint approximation. We use the precomputed LD scores of European individuals in the 1000 Genomes project for high-quality HapMap3 SNPs (‘eur_w_ld_chr’). For the phenotypic correlation, we consider 70 phenotypes excluding M79.6 (Enthesopathy of lower limb) and M67.8 (Other specified disorders of synovium and tendon), since the heritabilities of these two phenotypes estimated by LDSC are out of bounds. For the genetic correlation, we only consider 52 phenotypes exlcuding 20 phenotypes, where the heritabilities of these phenotypes are not significantly different from zero. We apply the K-means hierarchical clustering method to compare the correlations of phenotypes obtained by our proposed GPN and LDSC.

### Post-GWAS Analyses

#### Pathway enrichment analysis

To better understand the biological functions behind the SNPs identified by one multiple phenotype association test, we identify the pathways in which the identified SNPs are involved. We use the functional annotation tool named Database for Annotation, Visualization, and Integrated Discovery bioinformatics resource (DAVID: https://david.ncifcrf.gov/) (Huang et al. 2009; Sherman and Lempicki 2009) for the Kyoto Encyclopedia of Genes and Genomes (KEGG) pathway enrichment analysis. A mapped gene used in the pathway enrichment analysis denotes the gene that includes at least one identified SNPs with a 20kb window region. The biological pathways with FDR < 0.05 and enriched gene count > 2 are considered statistically significant (Cao et al. 2022).

#### Tissue enrichment analysis

To prioritize and interpret the GWAS signals and identify lead SNPs, tissue enrichment analyses are performed using the Functional Mapping and Annotation (FUMA: https://fuma.ctglab.nl/) (Watanabe et al. 2017) platform and the GWAS signals from one multiple phenotype association test in N.O. and in NET, respectively. FUMA first performs a genic aggregation analysis of GWAS association signals to calculate gene-wise association signals using MAGMA, which is a commonly used generalized gene-set analysis of GWAS summary statistics (de Leeuw et al. 2015). Then, it subsequently tests whether tissues and cell types are enriched for expression of the genes with gene-wise association signals. For tissue enrichment analysis, we use 30 general tissue types in GTEx v8 reference set (https://gtexportal.org/home/).

#### Colocalization analysis

As most associated variants are noncoding, it is expected that they influence disease risk through altering gene expression or splicing (Mountjoy et al. 2021). The colocalization analysis is a way to identify the association of a GWAS SNP and a gene expression QTL that are colocalized. We perform colocalization analysis using the ‘coloc’ package in R (Hormozdiari et al. 2016), a Bayesian statistical methodology that tests pairwise colocalization of eQTLs with unique identified SNPs by ceCLC in NET and N.O. from the UK Biobank dataset. The SNP-gene associations in the Muscle Skeletal tissue are downloaded from GTEx v7. We use the default setting of the prior probabilities, *p*_1_ = *p*_2_ = 10^-4^ and *p*_12_ = 10^-5^, for a causal variant in an eQTL or a GWAS SNP and a shared causal variant between eQTL and GWAS SNP, respectively.

## Availability of data and materials

### Data

The UK Biobank data are accessed via https://www.ukbiobank.ac.uk/ (Sudlow et al. 2015).

The GWAS catalog summary data are accessed via https://www.ebi.ac.uk/gwas/.

The SNP-gene associations in the Muscle Skeletal tissue are downloaded via https://gtexportal.org/home/.

### Software

The software for the proposed method is publicly available at https://github.com/xueweic/GPN.

PLINK version 1.9 can be downloaded from https://www.cog-genomics.org/plink/1.9/ (Chang et al. 2015).

LDSC: the command line tool for estimateing heritability and genetic correlation from GWAS summary statistiscs can be downloaded from https://github.com/bulik/ldsc (Bulik-Sullivan et al. 2015).

FUMA: the platform that can be used to annotate, prioritize, visualize and interpret GWAS results can be found from https://fuma.ctglab.nl/ (Watanabe et al. 2017).

DAVID: the functional tool can be found from https://david.ncifcrf.gov/ (Huang et al. 2009; Sherman and Lempicki 2009).

Cytoscape: an open source software platform for visualizing complex networks which can be accessed via https://cytoscape.org/ (Shannon et al. 2003).

## Acknowledgments

Part of this research has been conducted using the UK Biobank Resource under application number 41722 and the NHGRI-EBI GWAS Catalog. The work was in part funded by the Michigan Technological University Health Research Institute Fellowship program, the Portage Health Foundation Graduate Assistantship, and Graduate Dean Awards Advisory Panel. High-Performance Computing Shared Facility (Superior) at Michigan Technological University was used in obtaining results presented in this publication. Some parts of this work used the Extreme Science and Engineering Discovery Environment (XSEDE), which is supported by National Science Foundation grant number ACI-1548562. Specifically, it used the Bridges-2 system, which is supported by NSF award number ACI-1445606, at the Pittsburgh Supercomputing Center (PSC).

## Authors’ contributions

Formal analysis and Methodology: XC, SZ, and QS; Data curation and Visualization: XC; Writing original draft: XC, SZ, and QS; Writing review and editing: XC, SZ, and QS.

## References

Aschard H, Vilhjálmsson BJ, Greliche N, Morange P-E, Trégouët D-A, Kraft P. 2014. Maximizing the power of principal-component analysis of correlated phenotypes in genome-wide association studies. The American Journal of Human Genetics 94: 662–676.

Aterido A, Cañete JD, Tornero J, Ferrándiz C, Pinto JA, Gratacós J, Queiró R, Montilla C, Torre-Alonso JC, Pérez-Venegas JJ. 2019. Genetic variation at the glycosaminoglycan metabolism pathway contributes to the risk of psoriatic arthritis but not psoriasis. Annals of the Rheumatic diseases 78: 355–364.

Barber MJ. 2007. Modularity and community detection in bipartite networks. Physical Review E 76: 066102.

Blondel VD, Guillaume J-L, Lambiotte R, Lefebvre E. 2008. Fast unfolding of communities in large networks. Journal of statistical mechanics: theory and experiment 2008: P10008.

Bulik-Sullivan B, Finucane HK, Anttila V, Gusev A, Day FR, Loh P-R, Duncan L, Perry JR, Patterson N, Robinson EB. 2015. An atlas of genetic correlations across human diseases and traits. Nature genetics 47: 1236.

Bush WS, Oetjens MT, Crawford DC. 2016. Unravelling the human genome–phenome relationship using phenome-wide association studies. Nature Reviews Genetics 17: 129–145.

Bycroft C, Freeman C, Petkova D, Band G, Elliott LT, Sharp K, Motyer A, Vukcevic D, Delaneau O, O’Connell J. 2017. Genome-wide genetic data on~ 500,000 UK Biobank participants. BioRxiv: 166298.

Bycroft C, Freeman C, Petkova D, Band G, Elliott LT, Sharp K, Motyer A, Vukcevic D, Delaneau O, O’Connell J. 2018. The UK Biobank resource with deep phenotyping and genomic data. Nature 562: 203–209.

Cao X, Liang X, Zhang S, Sha Q. 2022. Gene selection by incorporating genetic networks into case-control association studies. European Journal of Human Genetics: 1–8.

Chang CC, Chow CC, Tellier LC, Vattikuti S, Purcell SM, Lee JJ. 2015. Second-generation PLINK: rising to the challenge of larger and richer datasets. Gigascience 4: s13742-13015-10047-13748.

Chung SA, Brown EE, Williams AH, Ramos PS, Berthier CC, Bhangale T, Alarcon-Riquelme ME, Behrens TW, Criswell LA, Graham DC. 2014. Lupus nephritis susceptibility loci in women with systemic lupus erythematosus. Journal of the American Society of Nephrology 25: 2859–2870.

Clauset A, Newman ME, Moore C. 2004. Finding community structure in very large networks. Physical review E 70: 066111.

Cole DA, Maxwell SE, Arvey R, Salas E. 1994. How the power of MANOVA can both increase and decrease as a function of the intercorrelations among the dependent variables. Psychological bulletin 115: 465.

Cordero AIH, Gonzales NM, Parker CC, Sokolof G, Vandenbergh DJ, Cheng R, Abney M, Sko A, Douglas A, Palmer AA. 2019. Genome-wide associations reveal human-mouse genetic convergence and modifiers of myogenesis, CPNE1 and STC2. The American Journal of Human Genetics 105: 1222–1236.

de Leeuw CA, Mooij JM, Heskes T, Posthuma D. 2015. MAGMA: generalized gene-set analysis of GWAS data. PLoS computational biology 11: e1004219.

Denny JC, Bastarache L, Roden DM. 2016. Phenome-wide association studies as a tool to advance precision medicine. Annual review of genomics and human genetics 17: 353–373.

Dey R, Schmidt EM, Abecasis GR, Lee S. 2017. A fast and accurate algorithm to test for binary phenotypes and its application to PheWAS. The American Journal of Human Genetics 101: 37–49.

Fine RS, Pers TH, Amariuta T, Raychaudhuri S, Hirschhorn JN. 2019. Benchmarker: an unbiased, association-data-driven strategy to evaluate gene prioritization algorithms. The American Journal of Human Genetics 104: 1025–1039.

Fortunato S, Barthelemy M. 2007. Resolution limit in community detection. Proceedings of the national academy of sciences 104: 36–41.

Fortunato S, Hric D. 2016. Community detection in networks: A user guide. Physics reports 659: 1–44.

Gaynor SM, Fagny M, Lin X, Platig J, Quackenbush J. 2022. Connectivity in eQTL networks dictates reproducibility and genomic properties. Cell Reports Methods 2: 100218.

Goh K-I, Cusick ME, Valle D, Childs B, Vidal M, Barabási A-L. 2007. The human disease network. Proceedings of the National Academy of Sciences 104: 8685–8690.

Gorlova O, Martin J-E, Rueda B, Koeleman BP, Ying J, Teruel M, Diaz-Gallo L-M, Broen JC, Vonk MC, Simeon CP. 2011. Identification of novel genetic markers associated with clinical phenotypes of systemic sclerosis through a genome-wide association strategy. PLoS Genet 7: e1002178.

Hawkins RD, Hon GC, Ren B. 2010. Next-generation genomics: an integrative approach. Nature Reviews Genetics 11: 476–486.

Hormozdiari F, Van De Bunt M, Segre AV, Li X, Joo JWJ, Bilow M, Sul JH, Sankararaman S, Pasaniuc B, Eskin E. 2016. Colocalization of GWAS and eQTL signals detects target genes. The American Journal of Human Genetics 99: 1245–1260.

Huang DW, Sherman BT, Lempicki RA. 2009. Bioinformatics enrichment tools: paths toward the comprehensive functional analysis of large gene lists. Nucleic acids research 37: 1–13.

Johnston KJ, Adams MJ, Nicholl BI, Ward J, Strawbridge RJ, Ferguson A, McIntosh AM, Bailey ME, Smith DJ. 2019. Genome-wide association study of multisite chronic pain in UK Biobank. PLoS genetics 15: e1008164.

Kim J, Bai Y, Pan W. 2015. An adaptive association test for multiple phenotypes with GWAS summary statistics. Genetic epidemiology 39: 651–663.

Kim SK, Nguyen C, Jones KB, Tashjian RZ. 2021. A Genome Wide Association Study For Shoulder Impingement and Rotator Cuff Disease. Journal of Shoulder and Elbow Surgery.

Kohane IS. 2011. Using electronic health records to drive discovery in disease genomics. Nature Reviews Genetics 12: 417–428.

Laird NM, Ware JH. 1982. Random-effects models for longitudinal data. Biometrics: 963–974.

Lee CH, Shi H, Pasaniuc B, Eskin E, Han B. 2021. PLEIO: a method to map and interpret pleiotropic loci with GWAS summary statistics. The American Journal of Human Genetics 108: 36–48.

Li R, Duan R, Kember RL, Rader DJ, Damrauer SM, Moore JH, Chen Y. 2019. A regression framework to uncover pleiotropy in large-scale electronic health record data. Journal of the American Medical Informatics Association 26: 1083–1090.

Liang K-Y, Zeger SL. 1986. Longitudinal data analysis using generalized linear models. Biometrika 73: 13–22.

Liang X, Cao X, Sha Q, Zhang S. 2022. HCLC-FC: A novel statistical method for phenome-wide association studies. Plos one 17: e0276646.

Liang X, Wang Z, Sha Q, Zhang S. 2016. An adaptive Fisher’s combination method for joint analysis of multiple phenotypes in association studies. Scientific reports 6: 1–10.

Mountjoy E, Schmidt EM, Carmona M, Schwartzentruber J, Peat G, Miranda A, Fumis L, Hayhurst J, Buniello A, Karim MA. 2021. An open approach to systematically prioritize causal variants and genes at all published human GWAS trait-associated loci. Nature Genetics 53: 1527–1533.

Newman M. 2018. Networks. Oxford university press.

Newman ME. 2012. Communities, modules and large-scale structure in networks. Nature physics 8: 25–31.

Newman ME, Girvan M. 2004. Finding and evaluating community structure in networks. Physical review E 69: 026113.

Nguyen T, Tagett R, Diaz D, Draghici S. 2017. A novel approach for data integration and disease subtyping. Genome research 27: 2025–2039.

O’Brien PC. 1984. Procedures for comparing samples with multiple endpoints. Biometrics: 1079–1087.

O’Connor LJ, Price AL. 2018. Distinguishing genetic correlation from causation across 52 diseases and complex traits. Nature genetics 50: 1728–1734.

O’Reilly PF, Hoggart CJ, Pomyen Y, Calboli FC, Elliott P, Jarvelin M-R, Coin LJ. 2012. MultiPhen: joint model of multiple phenotypes can increase discovery in GWAS. PloS one 7: e34861.

Pasaniuc B, Price AL. 2017. Dissecting the genetics of complex traits using summary association statistics. Nature Reviews Genetics 18: 117.

Pendergrass SA, Brown-Gentry K, Dudek S, Frase A, Torstenson ES, Goodloe R, Ambite JL, Avery CL, Buyske S, Bůžková P. 2013. Phenome-wide association study (PheWAS) for detection of pleiotropy within the Population Architecture using Genomics and Epidemiology (PAGE) Network. PLoS Genet 9: e1003087.

Pendergrass SA, Crawford DC. 2019. Using electronic health records to generate phenotypes for research. Current protocols in human genetics 100: e80.

Pendergrass SA, Dudek SM, Crawford DC, Ritchie MD. 2012. Visually integrating and exploring high throughput phenome-wide association study (PheWAS) results using PheWAS-view. BioData mining 5: 1–11.

Price AL, Patterson NJ, Plenge RM, Weinblatt ME, Shadick NA, Reich D. 2006. Principal components analysis corrects for stratification in genome-wide association studies. Nature genetics 38: 904–909.

Renauer PA, Saruhan Direskeneli G, Coit P, Adler A, Aksu K, Keser G, Alibaz Oner F, Aydin SZ, Kamali S, Inanc M. 2015. Identification of susceptibility loci in IL6, RPS9/LILRB3, and an intergenic locus on chromosome 21q22 in Takayasu arteritis in a genome wide association study. Arthritis & rheumatology 67: 1361–1368.

Sha Q, Wang X, Wang X, Zhang S. 2012. Detecting association of rare and common variants by testing an optimally weighted combination of variants. Genetic epidemiology 36: 561–571.

Sha Q, Wang Z, Zhang X, Zhang S. 2019. A clustering linear combination approach to jointly analyze multiple phenotypes for GWAS. Bioinformatics 35: 1373–1379.

Sha Q, Zhang Z, Zhang S. 2011. Joint analysis for genome-wide association studies in family-based designs. PloS One 6: e21957.

Shannon P, Markiel A, Ozier O, Baliga NS, Wang JT, Ramage D, Amin N, Schwikowski B, Ideker T. 2003. Cytoscape: a software environment for integrated models of biomolecular interaction networks. Genome research 13: 2498–2504.

Sherman BT, Lempicki RA. 2009. Systematic and integrative analysis of large gene lists using DAVID bioinformatics resources. Nature protocols 4: 44.

Solovieff N, Cotsapas C, Lee PH, Purcell SM, Smoller JW. 2013. Pleiotropy in complex traits: challenges and strategies. Nature Reviews Genetics 14: 483–495.

Stephens M. 2013. A unified framework for association analysis with multiple related phenotypes. PloS one 8: e65245.

Sudlow C, Gallacher J, Allen N, Beral V, Burton P, Danesh J, Downey P, Elliott P, Green J, Landray M. 2015. UK biobank: an open access resource for identifying the causes of a wide range of complex diseases of middle and old age. Plos med 12: e1001779.

Tachmazidou I, Hatzikotoulas K, Southam L, Esparza-Gordillo J, Haberland V, Zheng J, Johnson T, Koprulu M, Zengini E, Steinberg J. 2019. Identification of new therapeutic targets for osteoarthritis through genome-wide analyses of UK Biobank data. Nature genetics 51: 230–236.

Tang CS, Ferreira MA. 2012. A gene-based test of association using canonical correlation analysis. Bioinformatics 28: 845–850.

Terao C, Yamada R, Ohmura K, Takahashi M, Kawaguchi T, Kochi Y, Group HDGW, Kokubo M, Diop G, Yukawa N. 2011. The human AIRE gene at chromosome 21q22 is a genetic determinant for the predisposition to rheumatoid arthritis in Japanese population. Human molecular genetics 20: 2680–2685.

Tripathi B, Parthasarathy S, Sinha H, Raman K, Ravindran B. 2019. Adapting community detection algorithms for disease module identification in heterogeneous biological networks. Frontiers in genetics 10: 164.

Verma A, Bang L, Miller JE, Zhang Y, Lee MTM, Zhang Y, Byrska-Bishop M, Carey DJ, Ritchie MD, Pendergrass SA. 2019. Human-disease phenotype map derived from PheWAS across 38,682 individuals. The American Journal of Human Genetics 104: 55–64.

Visscher PM, Wray NR, Zhang Q, Sklar P, McCarthy MI, Brown MA, Yang J. 2017. 10 years of GWAS discovery: biology, function, and translation. The American Journal of Human Genetics 101: 5–22.

Wang M, Zhang S, Sha Q. 2022. A computationally efficient clustering linear combination approach to jointly analyze multiple phenotypes for GWAS. PloS one 17: e0260911.

Wang Z, Sha Q, Zhang S. 2016. Joint analysis of multiple traits using” optimal” maximum heritability test. PloS one 11: e0150975.

Watanabe K, Taskesen E, Van Bochoven A, Posthuma D. 2017. Functional mapping and annotation of genetic associations with FUMA. Nature communications 8: 1–11.

Xie H, Cao X, Zhang S, Sha Q. 2023. Joint analysis of multiple phenotypes for extremely unbalanced case-control association studies. Genetic Epidemiology doi:https://doi.org/10.1002/gepi.22513.

Yang JJ, Li J, Williams LK, Buu A. 2016. An efficient genome-wide association test for multivariate phenotypes based on the Fisher combination function. BMC bioinformatics 17: 1–11.

Yang Q, Wang Y. 2012. Methods for analyzing multivariate phenotypes in genetic association studies. Journal of probability and statistics 2012.

Zhou X, Stephens M. 2014. Efficient multivariate linear mixed model algorithms for genome-wide association studies. Nature methods 11: 407–409.

